# Evaluation of Deep Learning Algorithms to Predict Multiple Dementia-Related Neuropathologies from Brain MRI, Clinical and Genetic Data

**DOI:** 10.1101/2025.08.27.672683

**Authors:** Tamoghna Chattopadhyay, Rudransh Kush, Pavithra Senthilkumar, Christopher Patterson, Conor Owens-Walton, Emma J. Gleave, Sophia I. Thomopoulos, Sterling C. Johnson, Elizabeth C. Mormino, Timothy J. Hohman, Paul M. Thompson

**Affiliations:** Imaging Genetics Center, Mark and Mary Stevens Neuroimaging and Informatics Institute, Keck School of Medicine, University of Southern California, Marina del Rey, CA, USA; Wisconsin Alzheimer’s Disease Research Center, University of Wisconsin School of Medicine and Public Health, Health Sciences Learning Center, Madison, WI, USA; Department of Neurology and Neurological Sciences, Stanford University School of Medicine, Palo Alto, CA, USA; Wu Tsai Neurosciences Institute, Stanford University School of Medicine, Cogen Facility, Stanford, CA, USA; Department of Neurology, Vanderbilt University Medical Center, Nashville, TN, USA

**Keywords:** Alzheimer’s Disease, Deep Learning, Neuropathology Prediction, Multimodality

## Abstract

Alzheimer’s disease and related dementias (ADRD) involve overlapping neurodegenerative and vascular pathologies—such as amyloid-β (Aβ), tau, cerebral amyloid angiopathy (CAA), TDP-43, and alpha-synuclein—that complicate diagnosis and treatment. While PET and CSF biomarkers are useful for detecting Aβ and tau, they are invasive, expensive, and not widely available. In contrast, magnetic resonance imaging (MRI) is non-invasive and widely accessible, offering an opportunity for pathology prediction when combined with deep learning. Most prior studies have focused on single-pathology detection, but there remains a need for models that can jointly predict multiple co-occurring pathologies. In this work, we evaluate deep learning models that integrate structural MRI with demographic, clinical, and genetic data to classify six autopsy-confirmed neuropathologies: Aβ, tau, CAA, TDP-43, hippocampal sclerosis, and dementia with Lewy bodies. We compare our hybrid deep learning model to AutoGluon, an automated machine learning framework. Our findings support the potential of multimodal AI to enable non-invasive, comprehensive neuropathological profiling in ADRD.

## I. Introduction

Alzheimer’s disease and related dementias (ADRD)—including Alzheimer’s disease (AD), Parkinson’s disease (PD), dementia with Lewy bodies (DLB), and cerebrovascular pathologies—affect over 55 million people worldwide, a number expected to rise to 153 million by 2050 [1]. The societal and economic burden is staggering, with global costs projected to increase from $1.3 trillion in 2019 to $14.5 trillion by 2050 [1]. ADRD arises from a complex, heterogeneous set of co-occurring neuropathological processes, including amyloid-β (Aβ) plaques [2,3], hyperphosphorylated tau tangles [4,5], cerebral amyloid angiopathy (CAA) [6], transactive response DNA-binding protein 43 (TDP-43) [7,8] inclusions, and alpha-synuclein aggregates; the latter are characteristic of DLB and PD. These molecular pathologies frequently co-exist in the aging brain. For instance, CAA and TDP-43 pathology are each found in over 50% of AD cases [9-11], and they significantly contribute to cognitive decline independent of Aβ and tau pathology. This pathological overlap limits the effectiveness of single-target therapeutics, such as anti-amyloid or anti-tau agents [12-14], and presents a major challenge for differential diagnosis, trial stratification, and individualized treatment planning. As a result, classification systems such as ATN (Amyloid, Tau, Neurodegeneration) [15,16] are expanding to include vascular, inflammatory, immune, and synuclein-related biomarkers [17,18]. While positron emission tomography (PET) and cerebrospinal fluid (CSF) assays remain gold standards for detecting Aβ and tau in research contexts, their clinical utility is limited due to high cost, invasiveness, and limited availability. Furthermore, PET imaging does not capture other key pathological features such as TDP-43, CAA, or synucleinopathies [19]. In contrast, magnetic resonance imaging (MRI) is non-invasive and widely available, and can capture structural and microstructural brain changes associated with disease progression. Recent studies show that artificial intelligence (AI) models, particularly deep learning techniques, can extract disease-relevant features from multimodal MRI [20,21]. Kim et al. [22] employed a 2.5-D convolutional neural network—which processes sets of 2D slices from 3D FDG-PET volumes—to predict Aβ positivity, achieving an accuracy of 0.75 and an area under the curve (AUC) of 0.86. Similarly, Son et al. [23] used 2D CNN architectures to classify amyloid-PET images. Chattopadhyay et al. [24,25] compared various deep learning algorithms to predict Aβ positivity from structural T1w MRI. Wang et al. [26] used a 3D ResNet architecture with 3D T1w MRI to achieve balanced accuracies of 0.844, 0.839, and 0.623 for AD, vascular dementia (VaD), and DLB, respectively, showing strong pathology-specific classification performance. Still, there remains a critical gap in developing models that can jointly infer a range of co-occurring neuropathologies from whole-brain MRI in combination with clinical and genetic data. Such multi-target, multi-modal models could enable non-invasive, *in vivo* prediction of individual pathological profiles—advancing precision diagnostics and paving the way for tailored therapeutic interventions.

In this study, we evaluated deep learning models for predicting multiple neuropathological substrates, including Aβ, tau, CAA, TDP-43, hippocampal sclerosis (HPSC), and DLB, using structural MRI alongside demographic, clinical, and genetic covariates. By leveraging autopsy-confirmed labels from a large, diverse cohort, we aimed to assess the feasibility and utility of deep multimodal fusion models for comprehensive neuropathological classification in ADRD. We compare our custom hybrid deep learning model – that integrates tabular and image data – with the AutoGluon framework, on the classification task. AutoGluon [27]–an automated machine learning framework–serves as a robust baseline for comparison.

## II. Imaging Data and Preprocessing

The primary dataset for our experiments was a subset of the National Alzheimer’s Coordinating Center dataset [28] (N=950; age: 76.01 ± 11.86, 390F/560M) with both in vivo MRI and post mortem neuropathology. As HPSC and TDP-43 labels were unevenly distributed across diagnostic groups (625 dementia, 204 cognitively normal, 121 MCI), we manually balanced training, validation and test sets in the ratio 80:10:10. We ensured that there was no overlap between training and test data subsets, and that the test dataset had only one scan per subject. 3D volumetric T1-weighted (T1w) brain MRI volumes were pre-processed using the following steps [29]: nonparametric intensity normalization (N4 bias field correction), ‘skull stripping’ for brain extraction, nonlinear registration to an in-house template [29] with 6 degrees of freedom and isometric voxel resampling to 2 mm. The pre-processed images were of size 91×109×91. The T1w images were scaled using min-max scaling to take values between 0 and 1.

## III. Deep Learning Architectures

We developed a hybrid ‘Y-shaped’ deep learning architecture [30] that integrates a 3D convolutional neural network (CNN) with a parallel artificial neural network (ANN) to jointly model imaging and non-imaging data for neuropathology classification (**Fig. 1**). The 3D CNN branch processes volumetric T1-weighted MRI, using a series of convolutional blocks with increasing filter sizes (32, 64, 128, 256), each followed by batch normalization and max pooling. The final convolutional layer applies average pooling to capture global spatial patterns while reducing feature dimensionality. In parallel, the ANN receives normalized tabular inputs — including age, sex, diagnosis, time from imaging to death and from death to autopsy, APOE3 and APOE4 genotype count — passing them through three fully connected layers (1024, 512, 64) with instance normalization and ReLU activations. Categorical variables are one-hot encoded before being input to the network. Outputs from the CNN and ANN branches are concatenated and fed into dense layers for the final prediction. The model is trained end-to-end using the Adam optimizer (learning rate = 0.001, batch size = 2) for up to 100 epochs, with early stopping based on validation loss. Model performance is evaluated using balanced accuracy and F1 score.

**Figure 1.**
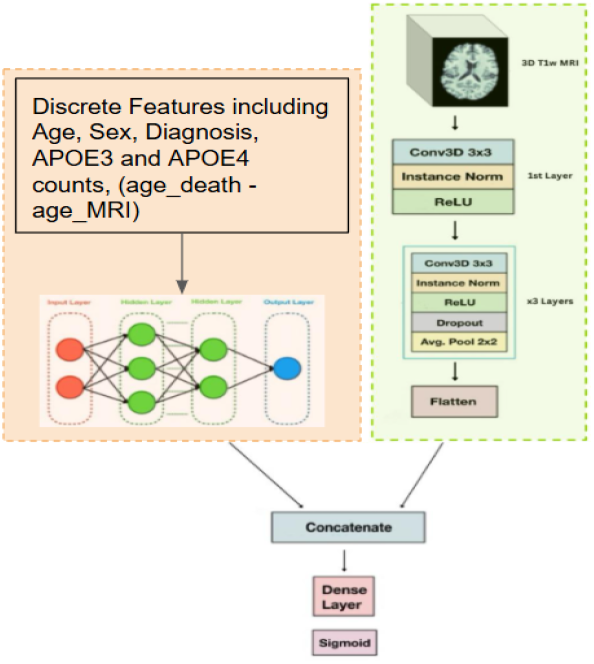
Hybrid 3D CNN Model Architecture. The Y-shaped design merges discrete features in tabular format (*left*) with entire 3D images (*right*), which are handled by a 3D CNN.

For comparison, we employed AutoGluon, an automated machine learning (AutoML) framework that can handle multimodal inputs, including tabular and image data, with minimal manual tuning (**Fig. 2**). Instead of using full 3D MRI volumes, we extracted the middle axial and coronal slices from each T1-weighted scan to reduce computational load while preserving key anatomical features.

**Figure 2.**
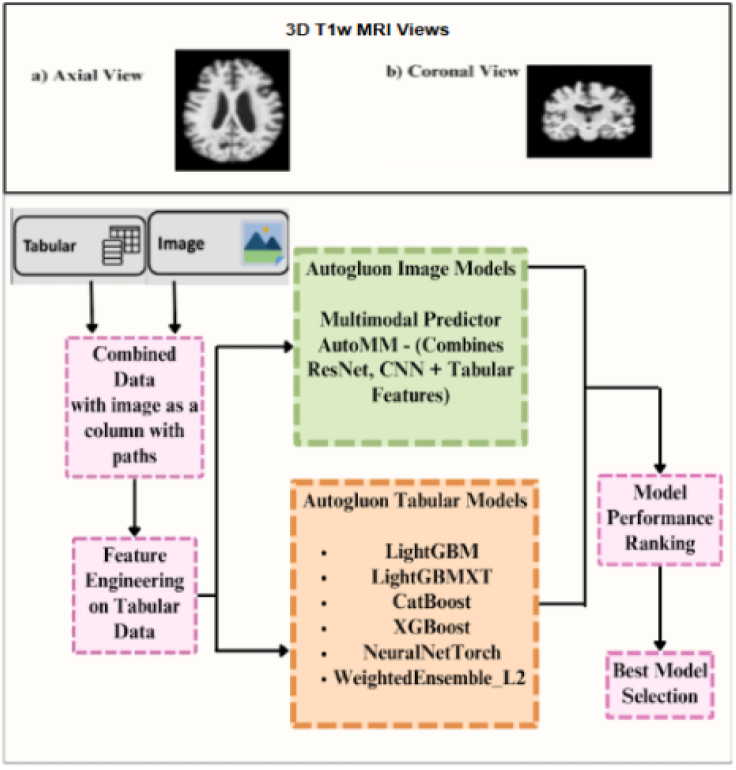
*Top:* Image inputs for the AutoGluon models. *Bottom:* AutoGluon architecture, which automatically selects a neural network architecture from a range of traditional models.

The same demographic and clinical variables used in our hybrid model were included as tabular input. AutoGluon automatically handled preprocessing, model selection, and hyperparameter tuning, and used ensemble strategies to improve generalization. Its performance was assessed using the same evaluation metrics as for our hybrid model. We evaluated multiple classification settings using our hybrid CNN framework. First, we trained individual binary classifiers for each neuropathological class: Aβ, tau, CAA, TDP-43, HPSC, and DLB. We then extended this to a multilabel classification task, enabling the model to simultaneously predict the presence or absence of multiple pathologies in each subject. In this set-up, a sigmoid activation function was applied to the output layer to produce independent probability scores for each pathology, capturing the non-mutually exclusive nature of neuropathological co-occurrence (in future, we could add a prior term with information on the empirical statistics of co-occurrence). We employed a binary cross-entropy loss function, summing across labels to guide optimization for each target independently within the multilabel setting. Label availability varied across the dataset—not all T1-weighted (T1w) MRI scans had ground truth annotations for every neuropathology—and we observed higher sample counts for amyloid-β, Lewy body disease, and HPSC, compared to CAA and TDP-43. Because of this, we conducted an additional multilabel classification experiment restricted to these three more frequently assessed pathologies. This allowed us to include a larger subset of the cohort, thereby increasing the effective training sample size (*N*) and potentially improving generalization. In a further extension, we incorporated tau pathology, using Braak staging as an additional label to form a six-pathology multilabel setting. For tau-specific analysis, we evaluated two strategies: a seven-class classification task using the full Braak scale (0–6), and a binary classification task, where Braak stages ≤3 were labeled as “low tau burden” and >3 as “high tau burden.” Due to architectural constraints, AutoGluon was only used for independent binary classification tasks, as it does not support multi-label outputs.

## IV. Results

The results, summarized in **Table 1**, show consistent trends across multiple experimental configurations. AutoGluon was first evaluated using 2D axial and coronal mid-slices extracted from 3D T1-weighted MRI (Cases 1 and 2). The model performed best for amyloid classification, but struggled to classify TDP-43 and hippocampal sclerosis (HPSC): F1 scores were 0.353 and 0.222, respectively. This may reflect the limited spatial context available in a single slice and the subtle or diffuse nature of these pathologies on structural MRI. No significant difference was observed between axial and coronal slices, suggesting that slice orientation had limited impact in this setting. In Case 3, we trained independent classifiers using the hybrid CNN. This configuration performed better across nearly all pathologies compared to AutoGluon. For instance, classification of TDP-43 reached a balanced accuracy of 0.719 and an F1 score of 0.690, while CAA achieved 0.763 and 0.711, respectively. The improved performance may stem from the model’s ability to leverage full volumetric spatial information and auxiliary demographic and clinical features.

**Table 1.** Performance of AutoGluon and Hybrid CNN models across various neuropathology classification tasks.

|  |  |  |  |  |  |  |  |  |
| --- | --- | --- | --- | --- | --- | --- | --- | --- |
|  |  | **TDP43** | **Amyloid** | **DLB** | **HPSC** | **CAA** | **Tau 2 Class** | **Tau Multi Class** |
| **Case 1 : AutoGluon Axial Slices** |  |  |  |  |  |  |  |  |
|  | **Balanced Accuracy** | 0.607 | 0.755 | 0.635 | 0.562 | 0.609 | 0.739 | - |
|  | **F1 Score** | 0.353 | 0.923 | 0.575 | 0.222 | 0.400 | **0.838** | - |
| **Case 2 : AutoGluon Coronal Slices** |  |  |  |  |  |  |  |  |
|  | **Balanced Accuracy** | 0.543 | 0.755 | 0.635 | 0.562 | 0.609 | 0.739 | - |
|  | **F1 Score** | 0.444 | 0.923 | 0.575 | 0.222 | 0.400 | **0.838** | - |
| **Case 3 : Hybrid CNN Individual Classifiers** |  |  |  |  |  |  |  |  |
|  | **Balanced Accuracy** | 0.719 | 0.744 | 0.671 | 0.669 | **0.763** | 0.758 | 0.735 |
|  | **F1 Score** | **0.609** | 0.695 | **0.683** | 0.255 | 0.711 | 0.831 | 0.463 |
| **Case 4 : Hybrid CNN MultiLabel Classifier for 3 Neuropaths** |  |  |  |  |  |  |  |  |
|  | **Balanced Accuracy** | - | **0.850** | 0.652 | 0.679 | - | - | - |
|  | **F1 Score** | - | **0.965** | 0.488 | 0.208 | - | - | - |
| **Case 5 : Hybrid CNN MultiLabel Classifier for all Neuropaths except Tau** |  |  |  |  |  |  |  |  |
|  | **Balanced Accuracy** | **0.735** | 0.740 | **0.690** | **0.732** | 0.710 | - | - |
|  | **F1 Score** | 0.520 | 0.901 | 0.603 | 0.308 | 0.733 | - | - |
| **Case 6 : Hybrid CNN MultiLabel Classifier for all Neuropaths and Tau** |  |  |  |  |  |  |  |  |
|  | **Balanced Accuracy** | 0.713 | 0.781 | 0.669 | 0.720 | 0.681 | **0.763** | - |
|  | **F1 Score** | 0.400 | 0.927 | 0.591 | **0.456** | **0.810** | 0.818 | - |

To assess the feasibility of multi-pathology classification, we evaluated the hybrid CNN in multilabel configurations. In Case 4, we restricted the model to the three most frequently assessed pathologies—amyloid, Lewy body pathology, and HPSC—to maximize sample size. While amyloid classification retained high accuracy, performance dropped for Lewy bodies (F1 score of 0.488) and HPSC (F1 score of 0.208), likely due to lower prevalence or subtler imaging features associated with those labels. In Case 5, we expanded the multilabel set-up to include CAA and TDP-43 in. While amyloid and CAA retained relatively high F1 scores, classification of Lewy bodies and TDP-43 became more challenging, perhaps due to increased label sparsity and overlapping phenotypic effects in the training data. In Case 6, we further included tau (based on Braak stage) in the multilabel classifier. The model maintained high performance for amyloid, tau, and CAA, suggesting that the fused multimodal architecture can effectively handle multiple co-occurring pathologies, even when trained jointly. Tau classification was also assessed independently in both binary and multiclass set-ups. The binary case consistently outperformed the multiclass set-up (7-way classification from stage 0 to 6). While the binary classifier achieved an F1 score of 0.831 in the individual model and 0.818 in the full multilabel setting, the multiclass task yielded lower performance (F1 score of 0.463), reflecting the challenge of differentiating adjacent Braak stages from MRI alone and potential sample imbalance across stages. Case 4 further highlights the trade-off between label completeness and model scope.

## V. Conclusions and Future Work

This study shows that it is feasible to use hybrid deep learning models to predict multiple neuropathological substrates from non-invasive MRI and clinical data, with superior performance over AutoML baselines. Our results underscore the potential of multimodal fusion approaches for advancing precision diagnostics in ADRD. Future work will focus on expanding to longitudinal MRI data, integrating additional imaging or biofluid modalities where available, and validating the models on external cohorts to ensure generalizability.

## Acknowledgments

This work was supported by NIH NIA grants U01 AG068057 (‘AI4AD’), U01AG082350 (‘CLARiTI’) and U24AG074855 (‘PHC’).

## Notes

### Competing Interest Statement

The authors have declared no competing interest.

